# Host- and microbial-mediated mucin degradation differentially shape *Pseudomonas aeruginosa* physiology and gene expression

**DOI:** 10.1101/2025.06.18.660398

**Authors:** Sabrina J. Arif, Kayla M. Hoffman, Jeffrey M. Flynn, Talia D. Wiggen, Sarah K. Lucas, Alex R. Villarreal, Adam J. Gilbertsen, Jordan M. Dunitz, Ryan C. Hunter

**Author notes:** To whom correspondence should be addressed: Ryan C. Hunter, Department of Microbiology & Immunology, Jacobs School of Medicine and Biomedical Sciences, University at Buffalo, 955 Main St., Buffalo, NY 14203. These authors contributed equally to this work (Sabrina J. Arif, Kayla M. Hoffman).

## Abstract

*Pseudomonas aeruginosa* is a hallmark pathogen of cystic fibrosis (CF) airway infections, capable of reaching high cell densities despite its limited ability to directly utilize mucin glycoproteins as a nutrient source. In the CF lung, however, *P. aeruginosa* may access preferred carbon sources (e.g., amino acids and short-chain fatty acids) through metabolic cross-feeding with co-colonizing mucin-degrading microbes. Although host-derived enzymes such as neutrophil elastase can also degrade mucins, the extent to which host-mediated mucin breakdown supports *P. aeruginosa* growth remains unclear. Thus, here we compared the nutritional impact of microbial versus host mucolytic activity on *P. aeruginosa* physiology. Analyses of CF sputum revealed patient-specific variability in mucin integrity that is shaped by both host and microbial factors. We demonstrate that mucin degradation by anaerobic bacteria through proteolysis, glycolysis, and fermentation, promotes robust *P. aeruginosa* growth, unlike mucin processed by neutrophil elastase alone. Targeted metabolomics identified acetate and propionate as key metabolites driving this cross-feeding, while transcriptomic and phenotypic analyses revealed that *P. aeruginosa* engages in diauxic growth on a broader set of mucin-derived substrates. Unexpectedly, cross-feeding with anaerobes triggered the induction of *P. aeruginosa* denitrification and fermentation pathways, suggesting redox remodeling despite being cultured under oxygen-replete conditions. Finally, the transcriptional profile of *P. aeruginosa* grown on anaerobe-conditioned mucins more closely resembled its in vivo gene expression, more so than when grown on intact or neutrophil-degraded mucins. Together, these findings provide new insight into the potential role of interspecies metabolic interactions in shaping pathogen physiology in the inflammatory, polymicrobial, and mucus-rich environment of the CF airways.

**Author Summary:** Cystic fibrosis (CF) airways contain viscous mucus that traps both pathogens and commensals. The major pathogen, *P. aeruginosa* thrives in these mucus-rich, inflamed environments, but how it acquires nutrients to sustain growth is poorly understood. We demonstrate that while host neutrophil enzymes degrade mucin polymers, this degradation alone does not provide substantial nutrients to support *P. aeruginosa* proliferation. In contrast, co-colonizing anaerobic microbiota extensively degrade mucins and generate short-chain fatty acids and other metabolites that strongly promote *P. aeruginosa* growth. We show that anaerobe-degraded mucins not only support faster growth but also trigger redox remodeling and gene expression changes that closely resemble *P. aeruginosa* behavior in CF patient sputum. This work highlights the important role of interspecies metabolic interactions in shaping CF airway infections and suggests new consideration for therapeutic strategies targeting airway microbiomes.

## Introduction

The gel forming mucins MUC5AC and MUC5B comprise approximately 97% of secreted airway mucins and serve as the structural backbone of the viscoelastic mucus barrier [1, 2]. In healthy airways, mucins are integral components of innate immunity, contributing to mucus rheology, sequestration of antimicrobials, and defense against inhaled pathogens [1, 3–8]. Conversely, mucin hypersecretion, aberrant glycosylation, proteolytic degradation, and changes in mucus viscosity and elasticity are common and are associated with increased susceptibility to persistent infection [9–14].

In the cystic fibrosis (CF) airways, mucins exhibit lower molecular weights compared to those from non-CF individuals, a phenotype thought to result from proteolytic degradation by neutrophil-derived enzymes [4, 9, 14–16]. CF sputum and bronchoalveolar lavage fluid (BALF) are enriched in neutrophils and their associated serine proteases, particularly neutrophil elastase, which is present in millimolar concentrations in the airway surface liquid [17–20]. NE has been shown to cleave mucins *in vitro,* preferentially targeting neutral, non-aromatic peptides [16], and in vivo evidence suggests host proteases degrade gel forming mucins during transport from the peripheral to central airways [21].

Although much of this proteolysis is attributed to inflammation [22], members of the airway microbiota also contribute to mucin degradation [22–24]. *Pseudomonas aeruginosa*, for example, produces several secreted metalloproteases, including LasA, elastase B, and alkaline protease, which are abundant in CF sputum [25–27]. Elastase B, in particular, can degrade mucins as effectively as NE [16, 26], and clinical isolates deficient in elastase production lack this capacity [28]. Mucins in sputum from chronically infected people with CF (pwCF) degrade more rapidly *ex vivo* than those from healthy controls, likely reflecting microbial mucolytic activity [14, 16]. Despite this, *P. aeruginosa* exhibits poor growth on intact mucin as a sole carbon source, suggesting that degradation alone is insufficient to support robust proliferation [29, 30].

Beyond canonical CF pathogens, the airways harbor a diverse community of strict and facultative anaerobic bacteria that are often overlooked in clinical diagnostics [29, 31, 32]. Many of these anaerobes possess potent mucin-degrading capabilities, similar to those found in the oral cavity and gastrointestinal tract. We previously showed that such activity is retained in CF sputum, where anaerobes contribute to mucin breakdown through proteolytic and glycolytic processes, liberating amino acids and glycans while generating short-chain fatty acids (SCFAs) such as acetate and propionate via mixed acid fermentation [29, 33]. These metabolites can enhance pathogen growth and virulence, and direct transcriptomic and metabolomic profiling of CF sputum supports the presence of in vivo cross-feeding interactions between anaerobes and *P. aeruginosa* [29].

Despite this evidence, it remains unclear whether host-derived mucin degradation, such as that mediated by NE, similarly expands the pool of bioavailable nutrients for *P. aeruginosa.* In this study, we compared the effects of host and microbial mucolytic processes on mucin integrity, nutrient availability, and *P. aeruginosa* physiology. We show that mucin degradation by anaerobes, but not NE, enhances *P. aeruginosa* growth, supporting a metabolic transition from glycolytic substrates to SCFAs (e.g., acetate), and promoting changes in cell morphology, redox state (NADH/NAD+ ratio), and core energy metabolism. Notably, anaerobe-conditioned mucins triggered the unexpected induction of denitrification and fermentation pathways, consistent with adaptation to a hypoxic, nutrient-rich environment. Finally, using a transcriptional accuracy scoring approach [34,35], we demonstrate that the gene expression profile of *P. aeruginosa* grown on anaerobe-degraded mucins more closely resembles its transcriptome in CF sputum, relative to growth on intact or NE-degraded mucins. These findings underscore the potential importance of interspecies metabolic interactions in shaping pathogen behavior and provide new insight into how *P. aeruginosa* may sustain infection within the mucus-rich, polymicrobial, inflammatory environment of the CF airways.

## Results

### Mucin integrity is variable in CF sputum and bronchoalveolar lavage

Given prior evidence of mucin degradation in the CF airways [27–30], we first assessed mucin integrity in expectorated sputum samples from 10 individuals with CF. Because no single method fully resolves mucin size, structure, and integrity [36], we applied complementary approaches to evaluate mucin degradation relative to intact controls; size exclusion chromatography (SEC) via fast protein liquid chromatography and ELISA-based detection of MUC5B.

Representative FPLC traces revealed two discrete peaks: (i) high molecular weight (HMW) mucins (>1000 kDa; peak 1), and (ii) lower molecular weight species (peak 2) **(Fig. 1A)**. Based on previously published studies [9] and our own analysis of FPLC molecular standards (**S1 Fig**), we interpreted the peak 1:peak 2 area-under-the-curve (AUC) ratio as a proxy for mucin integrity, with lower ratios indicating greater degradation. Purified salivary mucin (PSM) displayed high peak 1 AUCs, whereas CF sputum samples exhibited variable peak 1:peak 2 ratios (**Fig. 1B and S2 Fig**), suggesting inter-individual variability in mucin integrity.

**Fig. 1.**
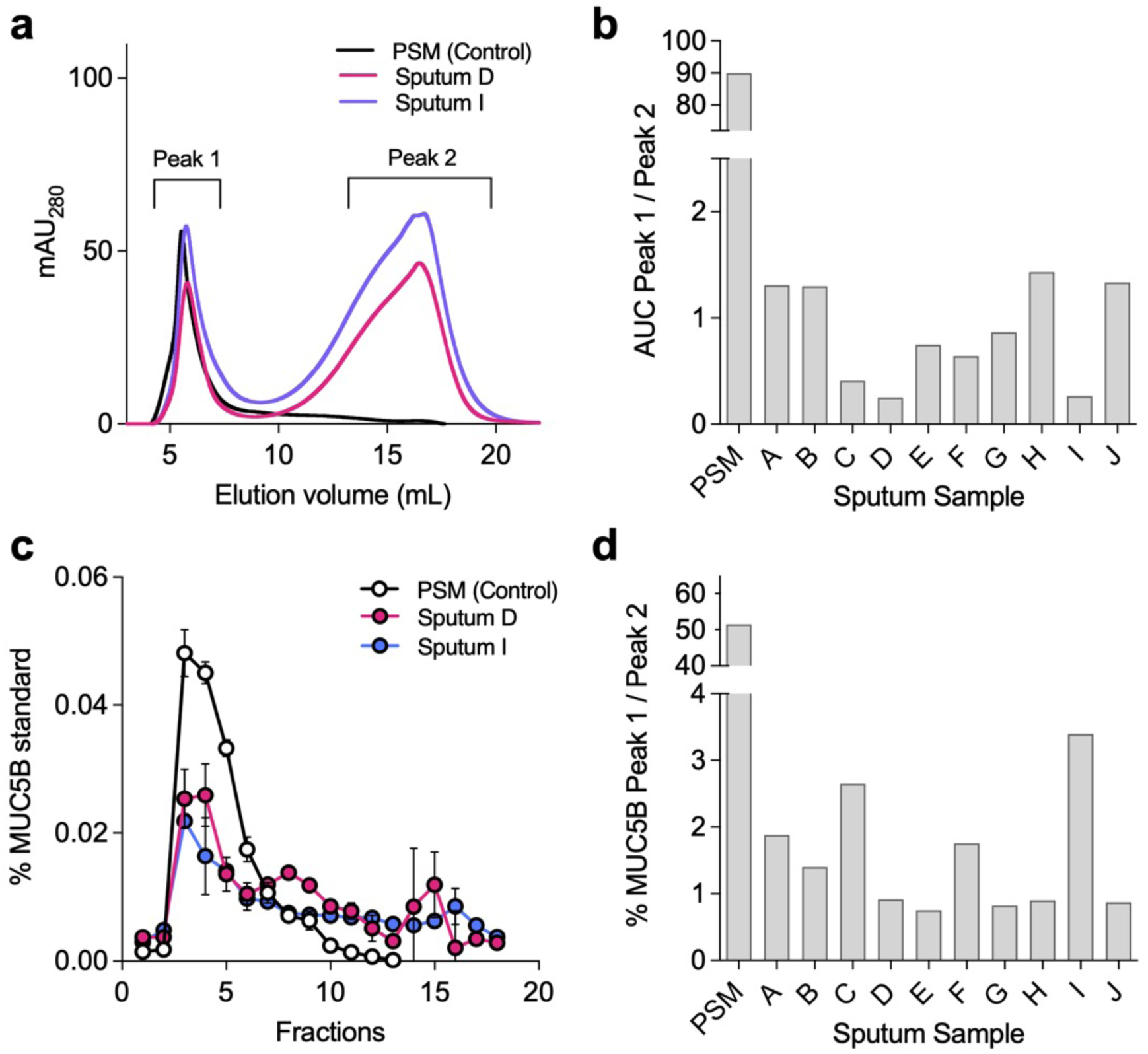
Mucin integrity is compromised in CF sputum. **(A)** Representative size exclusion chromatography (SEC) profiles of mucins isolated from CF sputum relative to purified salivary mucin (PSM, control). **(B)** Peak 1:Peak 2 area-under-curve (AUC) ratios from individual CF sputum samples (subjects A-J). **(C)** ELISA quantification of MUC5B immunoreactivity across SEC fractions. **(d)** Peak 1: Peak 2 ratios for MUC5B ELISA data. Error bars represent mean +/- standard deviation of triplicate measurements.

To evaluate whether degradation extended to the mucin polypeptide, SEC fractions were analyzed by ELISA for MUC5B immunoreactivity (**Fig. 1C,D**). As expected, intact mucins displayed MUC5B epitope reactivity concentrated in HMW fractions (fractions 2-5). In contrast, CF sputum exhibited lower immunoreactivity in peak 1 and detectable signal in peak 2 (fractions 9-15), consistent with proteolytic cleavage of the mucin backbone. Together, these data confirm substantial heterogeneity in mucin integrity across CF sputum samples.

### Both host and microbial enzymes contribute to mucin degradation

We next evaluated the relative contributions of host and microbial enzymes to mucin degradation. To do so, high molecular weight porcine gastric mucin (PGM), chosen for its commercial availability and ease of use, was purified and dialyzed to create a minimal mucin medium (MMM) as described previously [33]. MMM was then treated with either a physiologically relevant concentration (5 μg/mL) [19] of neutrophil elastase (‘NE’), an anaerobic mucin-degrading community (‘AMDC’, composed of *Veillonella, Fusobacterium, Prevotella spp.,* and other less abundant taxa (**S3 Fig**)), or both combined (AMDC+NE). Samples were incubated for 48h under anoxia and stored for further analyses.

Mucin integrity was then evaluated by FPLC as before (**Fig. 2A,B**). We note that due to preparation by the manufacturer, PGM integrity is comparatively low relative to purified human salivary mucin. However, when compared to untreated MMM, NE treatment alone shifted mucins from HMW to lower molecular weight fractions, as reflected by a reduction in peak 1 AUC and increase in peak 2 (**Fig. 2A,B)**. AMDC treatment similarly reduced peak 1 AUC, though to a lesser extent. Combined AMDC+NE treatment yielded additive degradation, producing the greatest loss of HMW material.

**Fig. 2.**
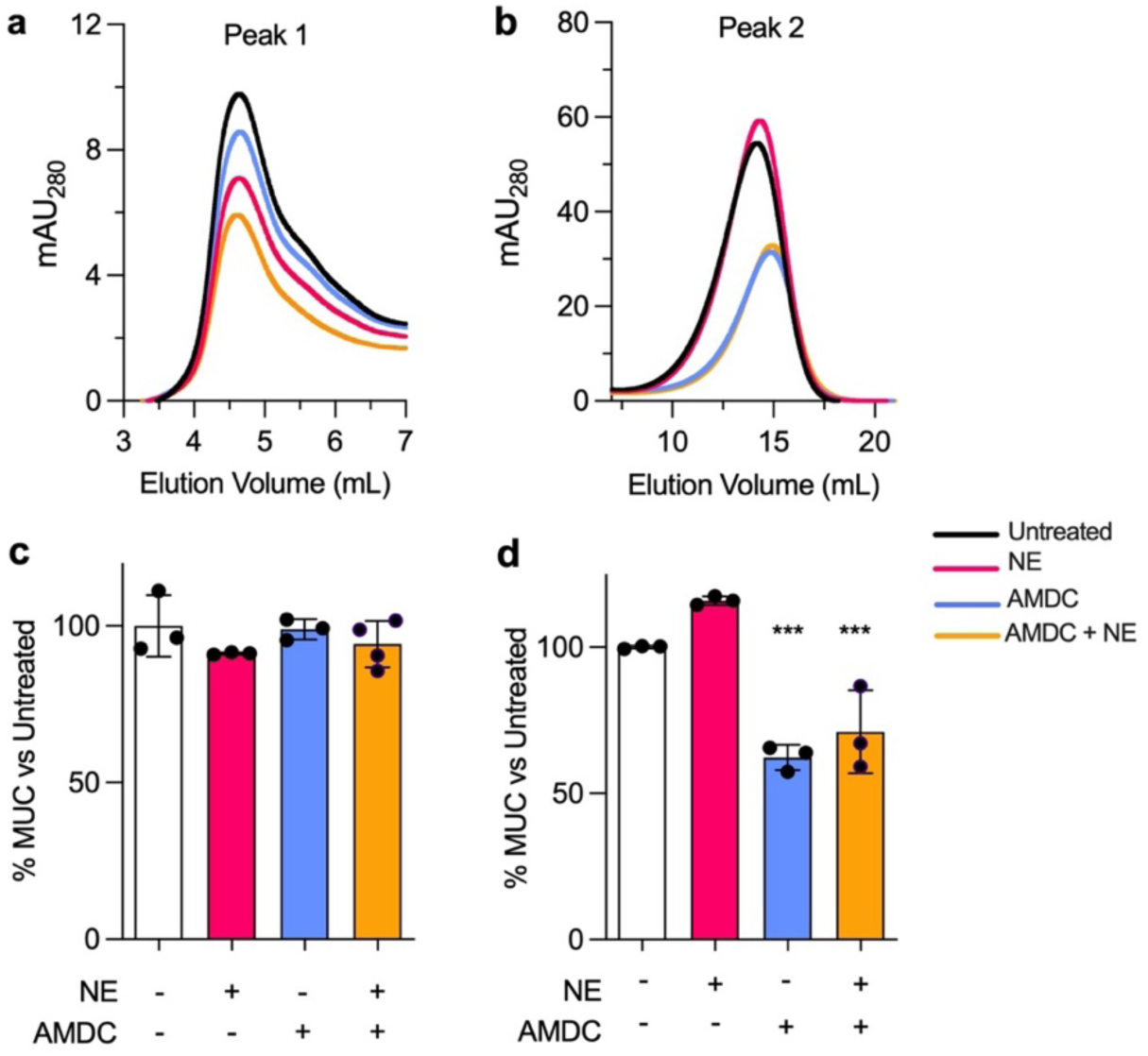
In vitro degradation by host and anaerobe mucinase activity. **(A,B)** FPLC-derived chromatograms of peak 1 (A) and peak 2 (B) after treatment with neutrophil elastase (NE), anaerobic mucin degrading community (AMDC), or both (AMDC+NE). **(C,D)** MUC5AC ELISA quantification of pooled peak 1 (C) and peak 2 (D) fractions. Error bars represent the mean and standard deviation of n=3 biological replicates with n=3 technical replicates each. ***, p<0.001.

Surprisingly, ELISA analyses showed that despite reductions in HMW material, MUC5AC immunoreactivity remained largely stable across treatments in peak 1 (**Fig. 2C**). In contrast, peak 2 immunoreactivity was markedly increased following anaerobic degradation (**Fig. 2D**), suggesting that anaerobes extensively process mucin polypeptides beyond the sites targeted by NE. These data demonstrate that anaerobic CF microbiota produce qualitatively distinct mucolytic effects compared to neutrophil elastase.

### Anaerobe-mediated mucin degradation enhances *P. aeruginosa* growth

We next tested whether mucin degradation products generated by host or microbial processes differentially supported *P. aeruginosa* growth. Growth media were prepared by normalizing mucin degradation supernatants (NE, AMDC, AMDC+NE) to equivalent total organic carbon (TOC) content relative to untreated minimal mucin medium (MMM). Under all conditions, *P. aeruginosa* exhibited similar early exponential growth (OD ∼0.2 at 10h) (**Fig. 3A)**. However, only AMDC- and AMDC+NE treated mucins yielded significantly higher final cell densities after 36h (p<0.001). In contrast, NE-treated mucin provided minimal additional growth benefit relative to untreated controls. Similar growth enhancement was observed across multiple clinical isolates, indicating that cross-feeding is not strain-specific (**Fig. 3B, S4 Fig**). These results indicate that anaerobe-mediated mucin degradation liberates nutrients not generated by NE alone and more effectively supports *P. aeruginosa* growth.

**Fig. 3.**
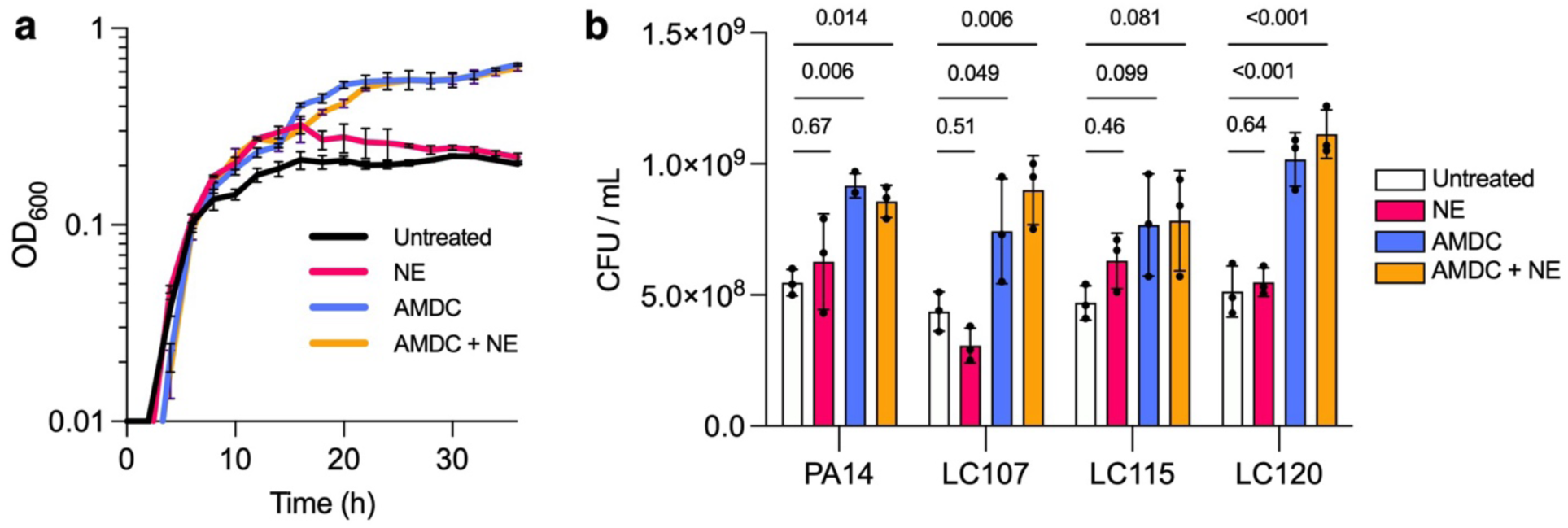
Anaerobic mucin degradation promotes *P. aeruginosa* growth. (**A**) Growth of *P. aeruginosa* PA14 on untreated, NE-treated, AMDC-treated, or AMDC+NE-treated mucins normalized by total organic carbon (TOC). (**B**) Final culture densities at 36h for PA14 and three CF clinical isolates. Full growth curves are shown in Fig. S4. Error bars represent mean +/- standard deviation and data shown are representative of three biological replicates for each experiment. Data were compared using a Kruskal-Wallis test with Dunn’s multiple comparisons.

### Short-chain fatty acids contribute to cross-feeding but do not fully explain enhanced growth

Given the abundance of SCFAs in CF sputum and BALF [29, 37, 38], we hypothesized that SCFAs generated by anaerobic fermentation drive the observed cross-feeding. HPLC analysis of mucin degradation supernatants confirmed that AMDC and AMDC+NE treatments produced substantial acetate and propionate, while NE treatment did not (as expected). All other metabolites tested (butyrate, lactate, succinate) fell below detection limits. During *P. aeruginosa* growth on AMDC-treated mucin, both acetate and propionate were progressively consumed, coinciding with increased cell density. Supplementing MMM with acetate and propionate reproduced much of the AMDC growth phenotype (**Fig. 4E**, compare black lines). However, *P. aeruginosa* mutants deficient in SCFA catabolic pathways ((Δ*aceA,* Δ*prpB,* Δ*acsA,* Δ*acsA*Δ*prpB*) retained the ability to grow on AMDC-conditioned mucins, suggesting that SCFAs alone do not fully account for the growth advantage. Other anaerobe-generated metabolites likely also contribute to nutrient availability.

**Fig. 4.**
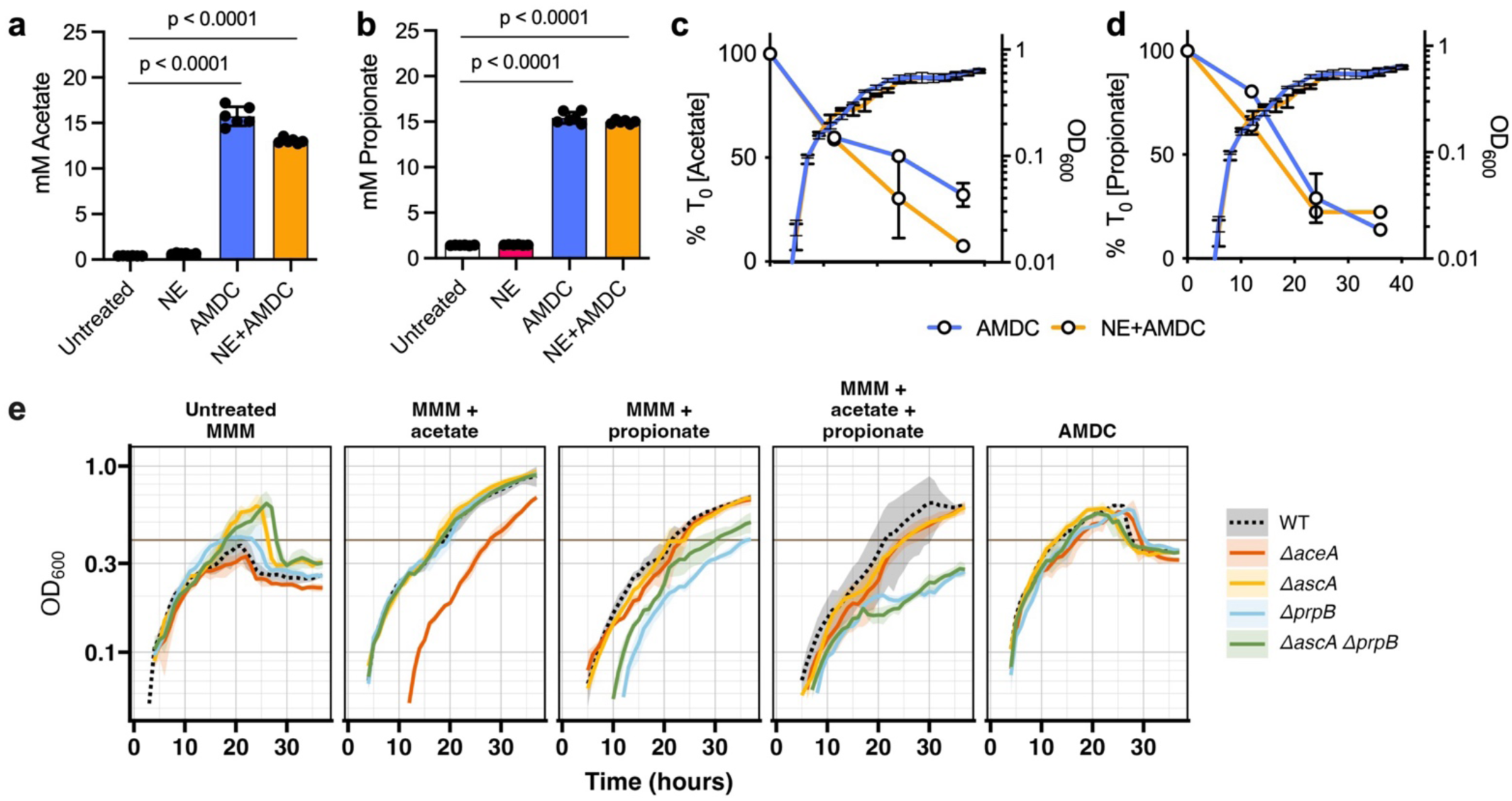
Short-chain fatty acid production and utilization by *P. aeruginosa.* **(A,B)** SCFA concentrations in mucin degradation supernatants. **(C,D)** Acetate and propionate depletion during *P. aeruginosa* growth (dashed lines). Change **(E)** Growth of WT PA14 and mutant strains (Δ*aceA*, Δ*acsA*, Δ*prpB*, and Δ*acsAΔprpB* double mutant on AMDC-treated on MMM supplemented with acetate, propionate, or both, and on AMDC-treated mucin. The dashed brown line indicates maximal OD for WT PA14 on untreated MMM. and AMDC-treated MMM supernatant. Maximal OD of wild-type *P. aeruginosa* PA14 grown on untreated MMM is denoted by the brown line (OD=∼0.4).

### Anaerobe-degraded mucins elicit distinct transcriptional responses in *P. aeruginosa*

To gain a deeper understanding of metabolites exchanged and the impact of anaerobe- and NE-mediated mucin degradation on *P. aeruginosa*, we performed RNAseq on *P. aeruginosa* grown on untreated, NE-treated, and AMDC-treated MMM. RNA was extracted after 4h and 16h, representing early to mid-log phase and the secondary mid-log phase observed in the AMDC condition, respectively. As anticipated, principal component analysis reflected growth phase (4h v. 16h), but at each time point, AMDC-treated cultures displayed distinct transcriptional profiles compared to NE-treated or untreated MMM (**Fig. 5A, B),** underscoring the discrete effect of anaerobe-derived metabolites on *P. aeruginosa* gene expression.

**Fig. 5.**
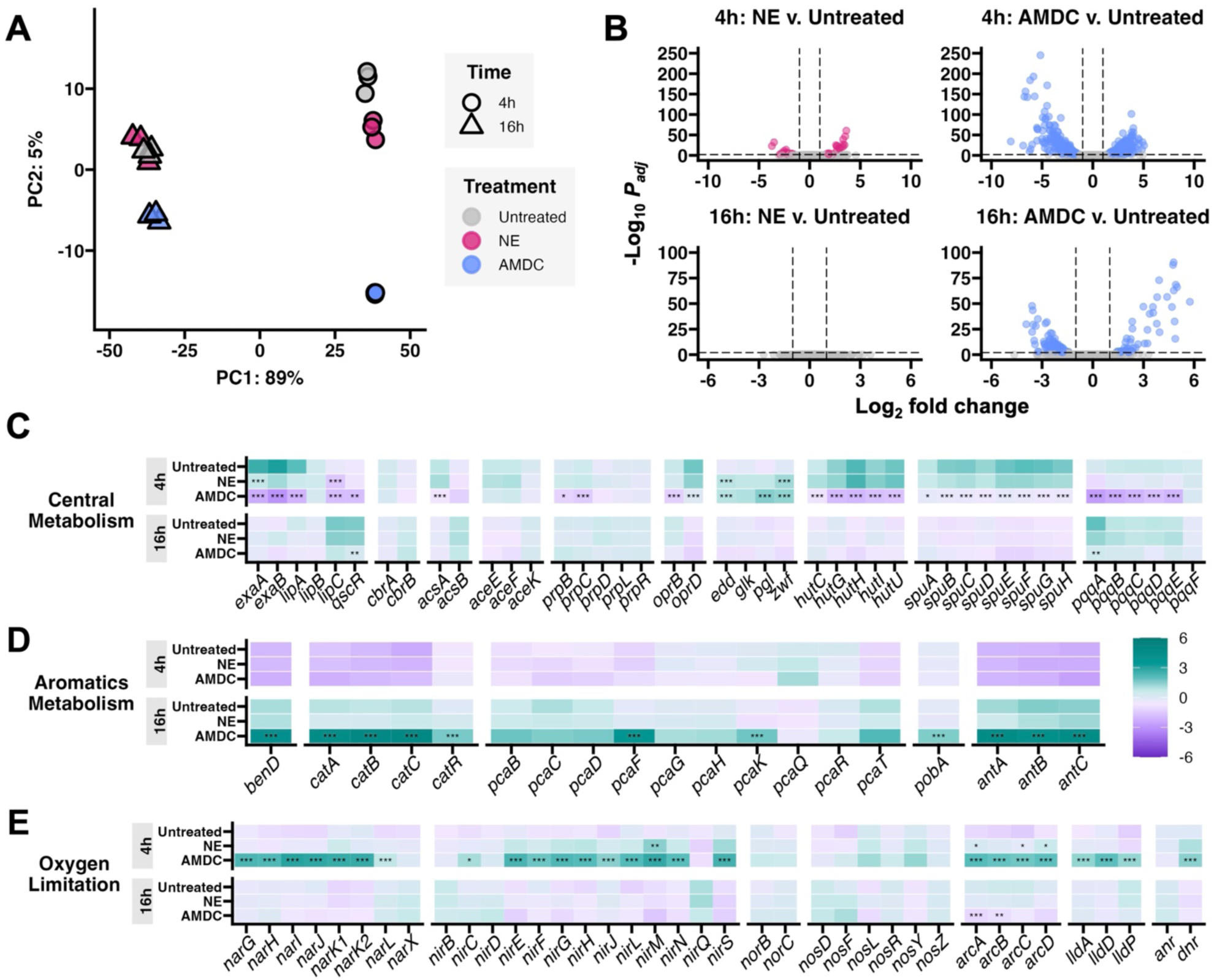
Transcriptomic remodeling during growth on anaerobe-degraded mucin. **(A) Principal component analysis of RNAseq data at 4h and 16h.missing description. (B)** Volcano plots of differential gene expression relative to untreated MMM. padj <0.01, log2fc>1. **(C,D,E)** Relative mean expression of genes involved in (**C)** central carbon metabolism, **(D)** aromatic metabolism, and (**E**) electron transport / redox pathways. Relative mean expression calculated as average gene expression among all conditions and timepoints. Significance: padj < 0.05, *; <0.01, **; or <0.001, ***.

Comparing untreated (MMM) and NE conditions revealed that at 4h, only 53 genes (30 upregulated, 23 downregulated) were differentially expressed (log_2_fold >1, padj<0.01)(**Fig. 5B, S1 File**). Upregulated genes were primarily associated with branched-chain amino acid degradation and Entner-Doudoroff/Embden-Meyerhof-Parnas pathways implicated in glucose metabolism. Downregulated genes included *lipC* (lipase), *exaA* (alcohol dehydrogenase), and the quorum sensing signal receptor gene, *qscR* (**Fig. 5C**). No genes were differentially expressed between untreated and NE culture conditions at 16h (**Fig. 5B, S1 File**), further supporting that NE-degraded mucins have limited effect on *P. aeruginosa* physiology.

Comparison of *P. aeruginosa* transcriptomes when grown on AMDC-degraded mucin relative to the untreated control revealed far greater differences in gene expression. At 4h and 16h, 530 genes (262 up, 268 down) and 195 genes (135 up, 60 down) were differentially expressed, respectively (**Fig. 5B, S1 File).** Many were associated with central metabolic pathways **(Fig. 5C)**, including *cbrB,* encoding a transcriptional regulator that increases expression of transporters and catabolic pathways in response to altered intracellular metabolite pools. Interestingly, despite depletion of acetate and propionate in the growth medium (see **Fig. 4B**), genes associated with acetate metabolism (*acsA*) and the glyoxylate shunt (*aceA, prpBC*) were unexpectedly downregulated during growth on AMDC-degraded mucin. We note, however, that this transcriptional pattern has been shown to occur in acetate-rich environments when other preferred carbon sources are also present [39]. Indeed, genes involved in glucose uptake (*oprB*) and its metabolism to glucose-6-phosphate (*glk*) were upregulated in AMDC-degraded medium at 4h, as were those involved in Entner-Doudoroff and pentose phosphate pathways (*zwf, pgl, edd,* and *eda*). Surprisingly, some genes (*zwf, edd*) were also upregulated in *P. aeruginosa* grown on NE-degraded mucins at 4h, which may suggest the catabolism of glucose or other mucin-derived monosaccharides during the early to mid-log phase of growth. Additionally, genes associated with the homogentisate pathway and other peripheral pathways were similarly upregulated at 4h, while downregulated pathways during early stages of growth included basic amino acid metabolism and transport (*hut, oprD*), polyamine utilization (*spu*), and pyrroloquinolone biosynthesis (*pqq)*(**Fig. 5C)**.

By 16h, genes involved in EDEMP pathways showed reduced expression in the AMDC condition, indicating a decreased reliance on glycolytic metabolism. Conversely, expression of genes responsible for the transport and catabolism of aromatic compounds, including the β-ketoapidate pathway (*ben, cat, pca, pob*), significantly increased. Interestingly, this pathway was previously shown to be upregulated in *Acinetobacter baumanii* due to mucin degradation [40]. Additionally, expression of *antA* (encoding anthranilate dioxygenase) increased, suggesting the metabolism of anthranilate, a precursor of tryptophan metabolism and the Pseudomonas quionolone signal (PQS). It is also worth noting that *ascA, aceA,* and *prpBC* showed increased expression at both time points. This suggests that while *P. aeruginosa* likely favors cleaved mucin glycans during early exponential growth, it may also use SCFAs as secondary carbon sources. This observation is consistent with their depletion from the growth medium (**Fig. 4B,D)** and suggests the likely exchange of additional metabolites.

### Anaerobe-derived metabolites drive redox remodeling despite oxygen-replete conditions

Despite oxygen-replete culture conditions, anaerobe-degraded mucin unexpectedly induced early activation of denitrification and fermentative pathways, including the *nar* and *nir* operons, encoding nitrate and nitrite reductases, respectively, in addition to arginine deiminase (*arcABCD*) and pyruvate fermentation (*lldH)* genes (**Fig. 5E).** These pathways, regulated by Dnr (an fnr-type transcription factor that was also highly expressed), are typically induced during hypoxia or redox imbalance. With the exception of *nirM,* upregulation of denitrification and fermentative pathways was not observed in the NE condition, suggesting that anaerobe-derived metabolites drive a remodeling of electron transport, among other pathways, in *P. aeruginosa*.

We next asked what advantage, if any, would be gained by *P. aeruginosa* in re-wiring redox pathways and electron transport machinery. One hypothesis is that increased expression of denitrification and fermentative pathways would enable *P. aeruginosa* to rapidly adapt to the dynamic oxygen conditions of the CF airways [41]. To test this possibility, *P. aeruginosa* was grown aerobically in MMM and ADMC-treated MMM for 4h, followed by a rapid shift to anoxic MMM containing 50mM nitrate. Contrary to our prediction, AMDC-“primed” *P. aeruginosa* did not gain any fitness benefit during the shift to anoxia (**Fig. 6A**), suggesting that anaerobic gene induction reflects redox rather than oxygen limitation per se.

**Fig. 6.**
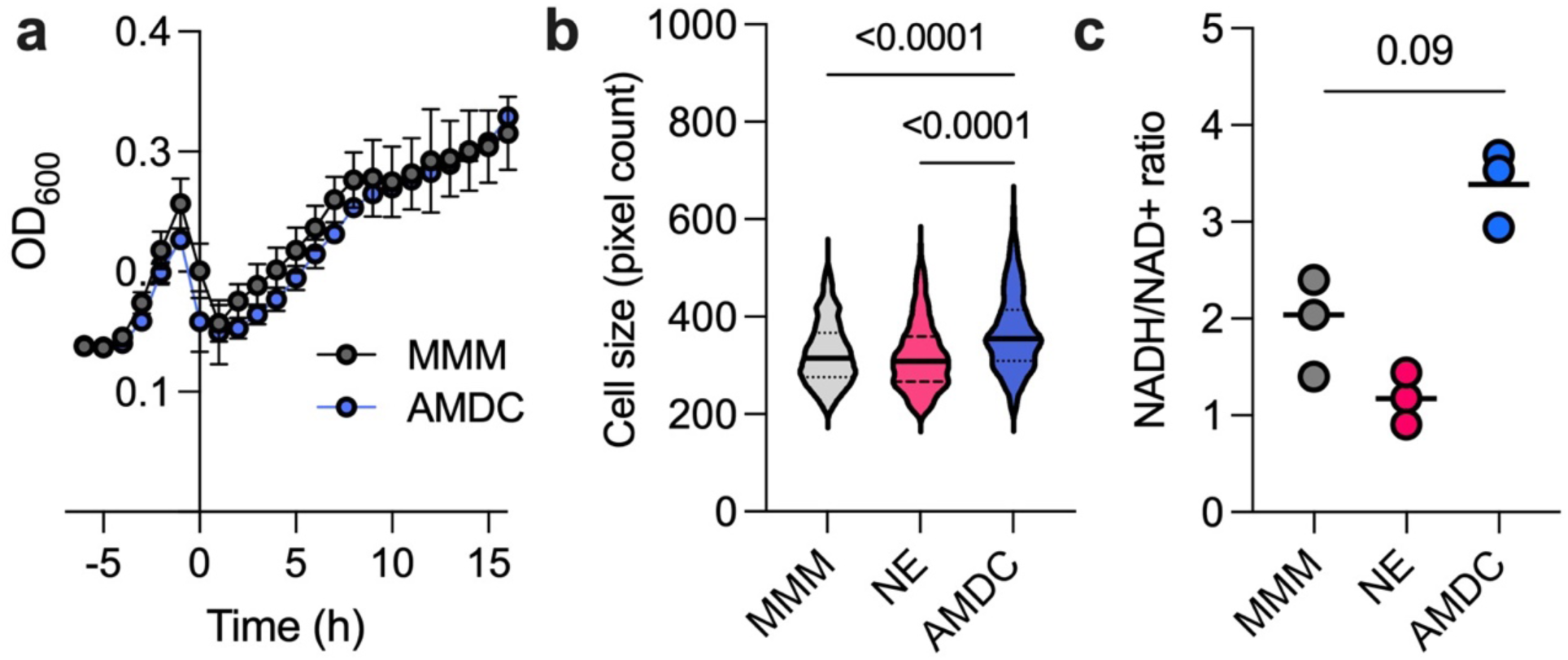
Anaerobe-degraded mucin alters *P. aeruginosa* redox state and cell morphology. (**A**) *P. aeruginosa* growth following oxic-to-anoxic shift (t=0) in minimal mucin medium (MMM) or AMDC-treated MMM (both media contained 50mM KNO_3_). Data shown are the mean +/ S.D. of three biological replicates. **(B)** cell size measurements after 4h aerobic growth. (**C**) Intracellular NADH/NAD+ ratio after 4h growth. Violin plots in (**B**) represent n=338 to 1533 bacterial cells captured across three images derived from three independent biological replicates. Data in (**C**) reflect the mean of three biological replicates. Data in (**B**) and (**C**) were compared using an ordinary one-way ANOVA with Tukey’s correction.

Consistent with this, *P. aeruginosa* grown on AMDC-treated mucin displayed larger cell sizes compared to cells grown on NE-degraded or untreated MMM (**Fig. 6B).** An increase in cell size is a recognized phenotypic response to excess electron donor availability [42,43]. Likewise, cells grown on AMDC-treated mucin exhibited a more reduced intracellular redox state, reflected by higher NADH/NAD+ ratios (**Fig. 6C)** relative to cells grown on NE-degraded or untreated MMM. Taken together, these results support a model in which rapid growth, fueled by anaerobe-derived metabolites, outpaces the respiratory capacity of the cell, triggering compensatory activation of less efficient denitrification and fermentation pathways to restore redox balance.

### *P. aeruginosa* gene expression on AMDC-degraded mucin is a better recapitulation of its *in vivo* transcriptional profile

Finally, to assess the *in vivo* relevance of the transcriptional changes observed, we applied an established accuracy score approach [34,35] that compares *P. aeruginosa* gene expression in vitro to transcriptomes derived from CF sputum. The AS2 metric calculates the proportion of genes whose expression *in vitro* falls within two standard deviations of their mean expression across 24 metatranscriptomes derived from CF sputum [34](**S2 File**). At 4h, *P. aeruginosa* grown on anaerobe-degraded mucin exhibited the highest median AS2 score (84.2%), compared to NE-treated mucin (81.5%) and untreated MMM (79.4%)(**Fig. 7A, S2 File**). These findings indicate that anaerobe-degraded mucins more closely approximate the in vivo transcriptional state of *P. aeruginosa* in CF airways. As expected, AS2 scores converged across conditions at 16h, consistent with earlier RNAseq data (**Fig. 5B)** showing reduced transcriptional divergence between conditions during later growth **(Fig. 7A)**.

**Fig 7.**
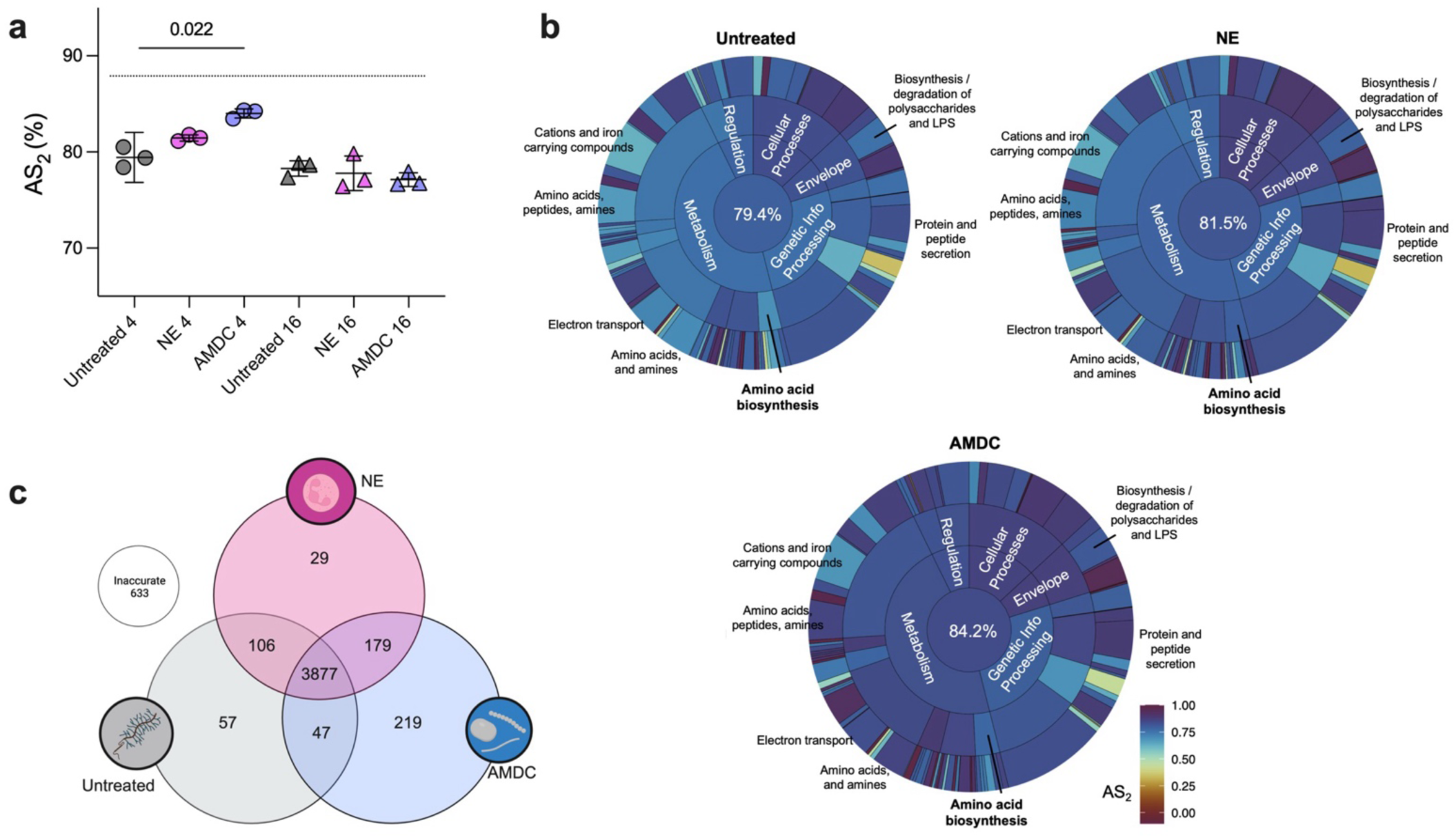
*P. aeruginosa* gene expression on anaerobe-degraded mucin better recapitulates CF sputum profiles. **A)** Accuracy score (AS2) comparison across conditions at 4h and 16h. Each data point represents a biological replicate (n=3 for each growth condition and time point). Dashed line represents the highest AS2 score obtained to date (Lewin et al., 2023). **(B)** Functional category accuracy (TIGRFAM meta, main, and sub roles). AS2 scores for all TIGRFAM categories are shown in S3 File. **(C)** Venn diagram of unique accurate genes across conditions. 633 genes were not accurate in any condition tested.

We next evaluated which specific functional pathways contributed to these differences. Using TIGRFAM functional annotations, we compared AS2 scores across metabolic, regulatory, and virulence-related gene categories, as previously described [34,35]. In nearly every major TIGRFAM functional role, AMDC-conditioned mucin either matched or exceeded untreated MMM in transcriptional accuracy (**Fig. 7B, S3 File**), with particularly strong improvements in metabolism, electron transport, protein secretion, iron homeostasis, and amino acid biosynthesis. NE-treated mucin modestly improved accuracy in a subset of categories but was consistently outperformed by AMDC across most functional pathways.

We also compared individual genes that were accurately expressed under each condition. AMDC- degraded mucin uniquely recapitulated the in vivo expression pattern for 219 genes, compared to 57 and 29 uniquely accurate genes in untreated and NE-treated mucins, respectively (**Fig. 7C**, **S4 File).** Many AMDC-specific accurate genes included those involved in redox metabolism (e.g. *narI, napC, nirC, cyoABDE*), acetate utilization (*acsA*), and denitrification pathways previously identified in our RNA-seq analyses (**Fig. 5C).** In contrast, some genes accurately expressed in untreated and NE conditions – such as multidrug efflux pumps (*mexB-oprM*), type VI secretion components (*vgrG4, tsi4*), and glyoxylate shunt enzymes (*aceA, glcB*) were less accurate in the AMDC condition, reflecting distinct metabolic adaptations. An additional 179 genes were accurate in both AMDC-treated and NE-treated mucin, but not in untreated MMM, suggesting that mucin degradation, regardless of the mucolytic source, can broadly influence *P. aeruginosa* gene expression.

## Discussion

*P. aeruginosa* is highly successful in establishing chronic infections in CF airways, yet the nutritional mechanisms that support its growth in vivo remain incompletely defined. While previous studies have demonstrated that *P. aeruginosa* can degrade mucins via its own proteases (e.g., LasB) and stimulate mucin production through host pathways such as EGFR signaling [27,30], its capacity to directly grow on intact mucins as a sole nutrient source is limited. In contrast, co-colonizing anaerobic bacteria, which are often enriched in CF airways, possess a broad repertoire of proteolytic and glycosidic enzymes that generate fermentation byproducts, particularly amino acids and SCFAs, that can serve as preferred carbon sources for *P. aeruginosa* [29,39]. However, the extent to which host versus microbial mucolytic activity shapes nutrient availability for *P. aeruginosa* has remained poorly defined.

Here, we systematically compared the effects of neutrophil elastase (NE; the dominant host protease in CF sputum) and mucin-degrading anaerobes on mucin integrity and *P. aeruginosa* physiology. While both NE and anaerobic bacteria degraded mucin polymers, only anaerobe-driven mucolysis significantly enhanced *P. aeruginosa* growth, even after normalizing for total carbon. NE-mediated degradation yielded smaller mucin fragments, but provided little additional nutritional value for the pathogen, highlighting that mucin proteolysis alone does not generate sufficient growth substrates.

In contrast, anaerobic degradation released abundant acetate and propionate which fueled robust *P. aeruginosa* growth. Interestingly, deletion of SCFA catabolism genes only partially impaired growth, suggesting that additional byproducts of microbial mucin degradation (e.g., amino acids, sugars, or other fermentation products) contribute to nutrient availability during cross-feeding. These results support a model in which anaerobic communities act as mucolytic “gatekeepers”, unlocking mucin-derived nutrient otherwise inaccessible to *P. aeruginosa*.

One unexpected finding was the induction of denitrification and fermentative pathways in *P. aeruginosa* grown on anaerobe-degraded mucin, even under fully oxygen-replete conditions. This metabolic remodeling is unlikely to reflect classical oxygen sensing, as anaerobe-primed cells did not exhibit enhanced growth following a shift to anoxia. Instead, we propose that rapid growth fueled by anaerobe-derived electron donors imposes redox pressure on *P. aeruginosa*, exceeding the capacity of aerobic respiration alone to reoxidize NADH. Under these conditions, cells increase in size, membrane space for respiratory complexes becomes limiting, and alternative electron acceptors (e.g., nitrate, pyruvate, arginine) are utilized to maintain intracellular redox balance [42,44]. Consistent with this model, *P. aeruginosa* grown on AMDC-degraded mucin displayed both increased cell size and elevated NADH/NAD+ ratios.

This form of metabolic flexibility may be highly advantageous in the CF lung, where steep oxygen gradients, fluctuating nutrient availability, and dense microbial communities create dynamic and spatially heterogeneous environments. Indeed, prior studies have documented denitrification pathway activation in CF sputum and airway mucus plugs, even when bulk oxygen is detectable [45,46]. Our data suggest that cross-feeding interactions with mucin-degrading anaerobes may contribute to this metabolic state by creating an electron donor-rich environment that functionally mimics aspects of hypoxia.

To quantitatively assess how well our in vitro models reflect in vivo behavior, we compared *P. aeruginosa* transcriptomes from mucin degradation conditions to CF sputum using the established accuracy score (AS2) framework [34,35]. AMDC-conditioned mucin yielded the highest transcriptional similarity to sputum samples, particularly during early growth, outperforming both NE-treated and untreated mucins. Functional improvements were observed across multiple metabolic pathways, including iron acquisition, electron transport, amino acid metabolism, and secretion systems – each a hallmark of *P. aeruginosa* adaptation during chronic CF infection. While synthetic sputum models remain superior in recapitulating the full complexity of chronic CF conditions [34,35,47], anaerobe-degraded mucin narrow this gap and may ultimately better approximate the metabolic state of *P. aeruginosa* encountered during earlier stages of disease, during their initial colonization stages when anaerobes are present at higher relative abundance.

Finally, while neutrophil processes are well-established contributors to mucin degradation in the CF airway [16,21], our data suggest that their role in shaping nutrient pools for *P. aeruginosa* is limited compared to that of mucin-degrading anaerobes. This distinction likely reflects fundamental differences in enzymatic activity: host proteases efficiently cleave polypeptide backbones but lack the broad glycosidase activities required to fully liberate sugars, SCFAs, and other fermentation intermediates. In contrast, anaerobic bacteria contribute a more expansive mucolytic repertoire, yielding a wider array of substrates that can support *P. aeruginosa* growth.

In summary, these findings highlight the critical role of interspecies metabolic interactions in shaping *P. aeruginosa* physiology within the mucus-rich, polymicrobial, and inflammatory environment of the CF airways. By elucidating how mucin degradation products generated by anaerobes fuel *P. aeruginosa* growth and alter its metabolic programs, data presented here provide new insight into the nutritional ecology of chronic CF infections.

## Methods

### Patient cohort and sample collection

Adult participants were recruited at the University of Minnesota Cystic Fibrosis Center. After informed consent, spontaneously expectorated sputum was collected, placed on ice, and transported to the UMN Department of Microbiology & Immunology for processing. Salivary mucins for control experiments were collected from healthy volunteers. This study was approved by the UMN Institutional Review Board (IRB#1511M80532).

### Mucin purification and isolation

High molecular weight mucins were purified from saliva for use as MUC5B and MUC5AC standards. To isolate mucin, saliva (∼50mL) was mixed with cOmplete protease inhibitor cocktail (Roche) and solubilized with 6M guanidine hydrochloride (GuHCl), 100mM Tris-HCl, 50mM dithiothreitol (DTT), and 4mM EDTA. Samples were incubated at 37°C for 1h, centrifuged at 15,000 rpm for 1hr, and dialyzed against MilliQ water at room temperature (2 x 1h) and 4°C (overnight). Lyophilized mucins were resuspended in 3M GuHCl, clarified by centrifugation (7000 rpm, 3 min), filtered (0.2μm), and purified by fast protein liquid chromatography (FPLC; ÄKTA Pure, Cytiva) using a 10/200mm Tricorn column packed with Sepharose CL-2B. Aliquots (500 μL) were loaded and eluted isocratically at 0.4mL/min in 50mM phosphate buffer (pH 7.2) containing 150mM NaCl. Fractions were frozen at −80°C for storage.

### Mucin analysis by size exclusion chromatography

For sputum samples, 6M GuHCl was added at a 6:1 (v/v) ratio, followed by incubation at 80°C for 2h with intermittent mixing. Samples were clarified using 0.45μm centrifugal filters (Thermo) and analyzed via FPLC as above. Eighteen 1-mL fractions were collected and stored at −80°C for downstream ELISA. Mucins derived from bacterial cultures were similarly processed after debris removal by centrifugation and filtration. Area-under-the-curve (AUC) calculations were performed using UNICORN 7.0 software (Cytiva).

### Enzyme linked immunosorbent assays (ELISA)

For each sample, fractions from peak 1 (fractions 2-5) and peak 2 (fractions 9-16) were pooled. Standards were generated using serial dilutions of purified salivary mucin. Samples and standards (50μL) were applied in triplicate to block MaxiSorp (Thermo), dried at 45°C (∼2.5 h) until liquid had evaporated, washed three times with 300μL TBS-T (1X TBS + 0.1% Tween 20) and blocked with 300μL Protein-Free Blocking Buffer (Thermo) for 1h at RT. For MUC5B detection, plates were incubated with anti-MUC5B antibody (8C11) (Sigma) diluted 1:1000 in TBS-T + 2% BSA for 1h at RT. Plates were washed three times TBS-T and incubated with goat-anti-mouse HRP (Thermo) diluted 1:2000 in TBS-T+ 2% BSA for 1h. For MUC5AC, anti-MUC5AC (45M1)(Thermo) was used, followed by HRP-conjugated secondary (1:1000). SuperSignal ELISA Femto Substrate (Thermo) was used for luminescence detection on a Synergy H1 plate reader (BioTek).

### Bacterial strains and growth conditions

*P. aeruginosa* strain PA14, three clinical isolates (LC107, LC115, LC120), and isogenic mutants of PA14 (*ΔaceA*, *ΔacsA*, *ΔprpB*, and *ΔacsΔprpB*) generated previously [29] were maintained on Luria Bertani (LB) broth or agar. The anaerobic mucin-degrading bacterial community (‘AMDC’, **S3 Fig**) was revived from freezer stock and passaged twice (48h) on a defined minimal mucin medium (see below) in an anaerobic chamber (95% N^2^, 5% H_2_, 5% CO_2_) (Coy Laboratory Products).

### *In vitro* mucin degradation

A minimal mucin medium (MMM) was prepared by dissolving porcine gastric mucin (PGM) (Sigma) in Milli-Q water (30g/L), autoclaved, and clarified as described previously [33]. The resulting MMM contained 50mM KH2PO4, 150mM NaCl, 1mM MgSO4, and a trace mineral mix [29]. MMM was incubated with (i) no treatment, (ii) AMDC (OD_600_ = 0.3), (iii) 5 µg/mL neutrophil elastase (Abcam), and (iv) both AMDC and NE together. All cultures were incubated anaerobically at 37°C for 48h. Degradation was assessed by FPLC and ELISA as described above.

### Total organic carbon (TOC) analysis

7mL volumes of *in vitro* mucin degradation samples were frozen and lyophilized as described above. 150mg of dehydrated sample were analyzed for TOC using a Vario Max Cube elemental analyzer (Elementar). TOC-normalized supernatants (normalized to 14% TOC) were used for subsequent *P. aeruginosa* growth assays.

### P. aeruginosa growth assays

*P. aeruginosa* was cultured overnight in LB and washed three times with phosphate buffered saline (PBS). 1 x 10^5^ CFU/mL were inoculated into 200µL of TOC-normalized mucin degradation supernatants in 96-well plates, sealed with Breathe Easy membranes (Diversified Biotech) and incubated aerobically at 37°C with orbital shaking in a Synergy H1 microplate reader (BioTek). OD_600_ was measured hourly for 36h. All conditions were performed with three biological and three technical replicates. *ΔaceA*, *ΔacsA*, *ΔprpB*, and *ΔacsΔprpB* mutant growth assays were similarly performed using MMM or AMDC-supernatants supplemented with 15mM acetate, 15mM propionate, or both. For oxic-to-anoxic shift experiments, PA14 was grown in MMM or AMDC-MMM aerobically for 4h, then transferred to an anaerobic chamber and inoculated 1:10 into MMM + 50mM KNO_3_. Anaerobic growth was monitored for 16h in microplate readers housed within the chamber.

### High performance liquid chromatography (HPLC)

Mucin degradation supernatants were clarified by centrifugation, filtered through a 3,000 MWCO PES concentrator (Thermo) and analyzed using a Dionex UltiMate 3000 HPLC (Thermo) equipped with an Acclaim organic acid column (4.0 x 250 mm, 5 μm). Samples (10μL) were eluted isocratically (100mM Na_2_SO_4_, pH 2.6 with CH_3_SO_3_H) at 1mL/min for 24h following an 8 min incubation step [48]. SCFAs were quantified relative to standard curves.

### RNA extraction and sequencing

PA14 cultures were grown for 4h or 16h in TOC-normalized mucin degradation supernatants (60mL each), harvested by centrifugation, and snap-frozen. RNA was extracted using the RNeasy Mini Kit (Qiagen) with β-mercaptoethanol lysis and DNaseI treatment, followed by cleanup with the RNA Clean and Concentrator kit (Zymo). rRNA depletion was performed using Ribo-Zero Plus [Illumina], and libraries were prepared using TruSeq RNA Library Prep kit (Illumina) per manufacturer’s instructions. Sequencing was performed on the Illumina NovaSeq6000 platform (50 bp paired-end reads) at the University of Minnesota Genomics Center.

### RNA-seq analysis

Raw fastq files were quality-checked (FastQC), aligned to the *P. aeruginosa* PA14 genome (NCBI RefSeq 008463.1) using the R ‘Subread’ package [49], and annotated via Pseudomonas.com. Features with low read counts (<10) or non-protein coding genes were filtered. DESeq2 [50] was used for normalization, principal component analysis, and differential gene expression testing (log_2_fold-change > 1, padj <0.001). Code and data are shared at https://github.com/Hunter-Lab-UMN/Arif_SJ_2025. Raw RNA-seq files are deposited under NCBI bioproject PRJNA1279252.

### Accuracy score analysis

Accuracy scores (AS2) were calculated as previously described [34, 35]. RNAseq data were realigned to the *P. aeruginosa* PA01 genome to enable comparison to published CF sputum metatranscriptomes. For each gene, z-scores were calculated relative to the mean expression across 24 CF sputum samples [34]. AS2 scores represent the percentage of genes falling within +/- 2 standard deviations of in vivo expression. TIGRFAM category-level AS2 values were similarly computed. Sunburst plots were generated using the ggsunburst R package [51]. Statistical comparisons were performed using Kruskal-Wallis tests with Dunn’s post-hoc correction in Prism 10.

## Acknowledgements

We thank current and former members of the Hunter lab for discussions and manuscript feedback, as well as the attendees of the Atlanta CF Models Workshop (Georgia Institute of Technology, 2023, supported by the Cystic Fibrosis Foundation, #WHITEL20A0), particularly Gina Lewin, Rebecca Duncan, and Marvin Whiteley for assistance with the accuracy score framework. We also thank Kristin Jesse and the University of Minnesota Cystic Fibrosis center for patient recruitment and sample acquisition.

