## Supplemental Data for "Host- and microbial-mediated mucin degradation differentially shape *Pseudomonas aeruginosa* physiology and gene expression"

**S.J. Arif, K.M. Hoffman et al.**

**Document S1.** Figures S1-S4

**S1 File.** *P. aeruginosa* differential gene expression in untreated (MMM), NE-treated, and AMDC-treated mucin.

**S2 File**. Raw counts, normalized sequence counts, and accuracy scores for the core *P. aeruginosa* gene set (5147 genes)

**S3 File.** AS2 and z-scores for TIGRFAM-annotated *P. aeruginosa* genes and their expression during growth on intact and degraded mucins.

**S4 File.** Individual gene expression accuracy

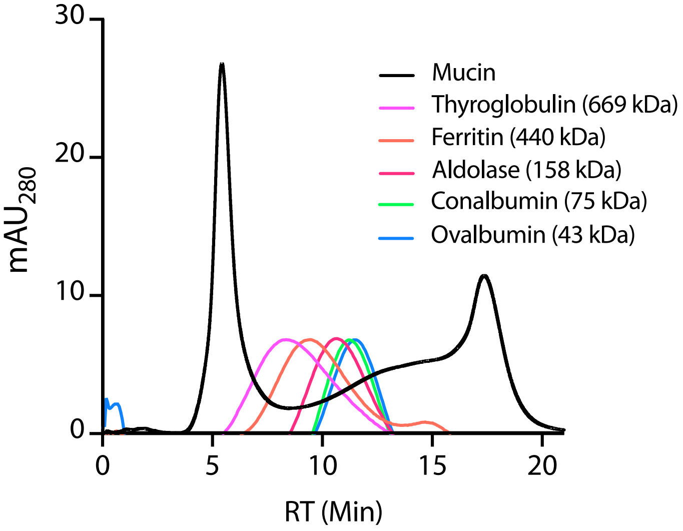


**S1 Fig.** High molecular weight standards in the GE Gel Filtration Calibration Kit as analyzed by FPLC. Higher molecular weight compounds (i.e. intact mucin) elute earlier than those of lower molecular weight (i.e. degraded mucin).

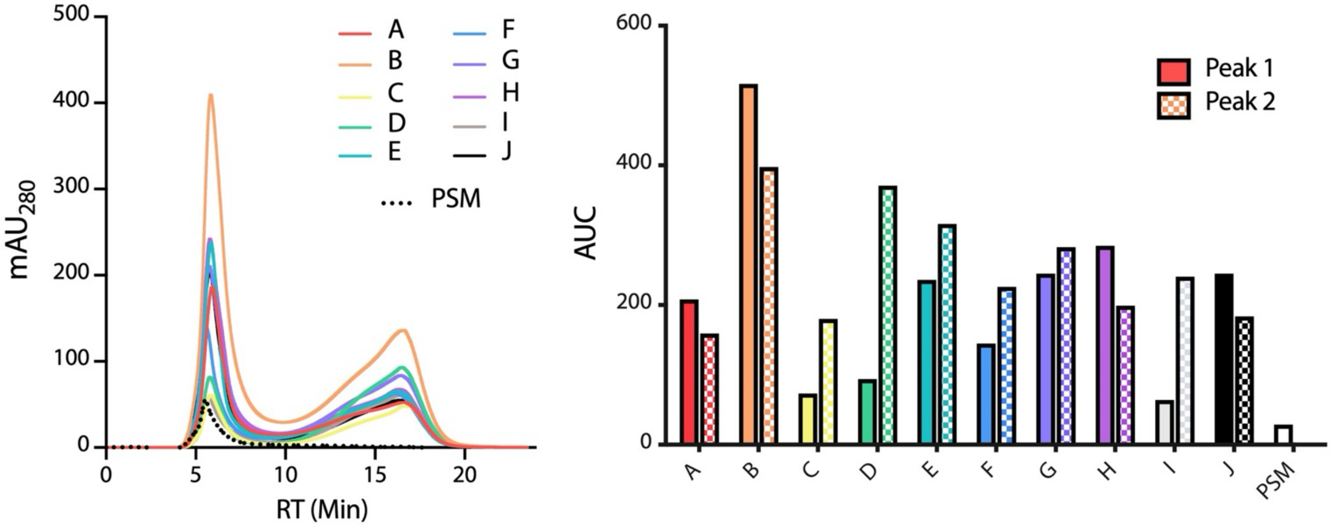


**S2 Fig. Mucin integrity in CF sputum**. **(a)** Size exclusion chromatography profiles of mucins isolated from CF sputum relative to purified sinus mucin (PSM, dashed line). **(b)** Area-under-curve (AUC) data for Peak 1 (solid color) and Peak 2 (hatched color) of individual sputum samples compared to PSM.

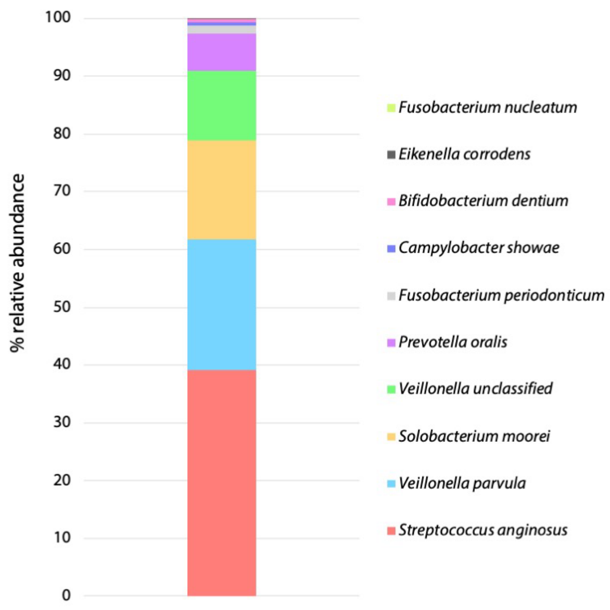


**S3 Fig.** Composition of the anaerobic mucin degrading community (AMDC) as determined by 16S rRNA gene sequencing.

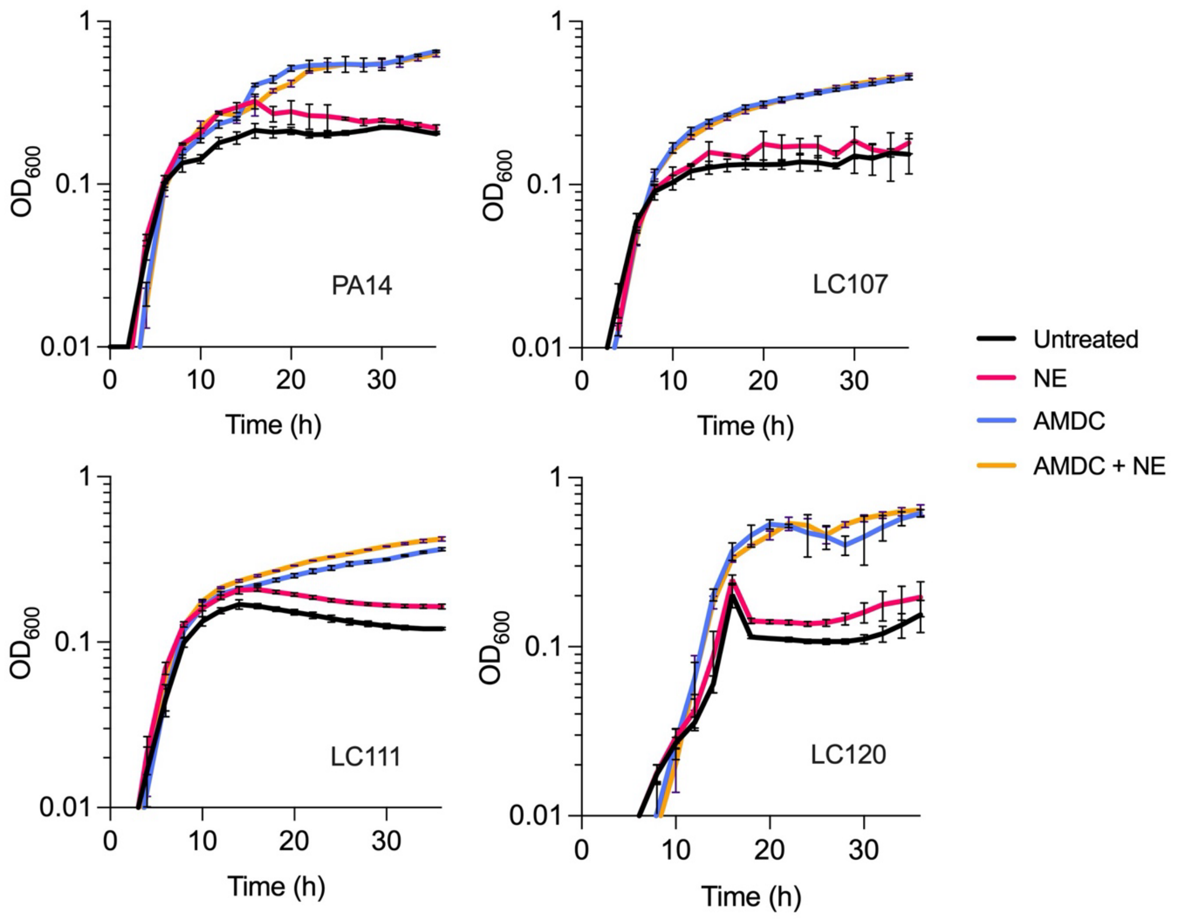


**S4 Fig.** Growth curves of *P. aeruginosa* clinical isolates on untreated MMM, and NE-, AMDC-, and AMDC+NE-treated MMM. Each isolate exhibited increased growth in the cell-free supernatants of AMDC- and AMDC+NE-treated mucin.
